# Extracts of *Bacillus proteolyticus* strain OSUB18 exhibit cytostatic activity

**DOI:** 10.64898/2026.09.30.755762

**Authors:** Baorui Kang, Dalton Hironaka, Pu Yuan, Darwin Munoz, Kou-San Ju, Ye Xia, Gaofeng Xiong

## Abstract

Although therapeutic advancements have dramatically improved outcomes for cancer patients in recent decades, advanced cancer remains a significant challenge. There is still an urgent need for new and more effective strategies that minimize toxicity and improve survival outcomes in cancer patients. Natural products from microorganisms are a rich and promising source of bioactive compounds with substantial therapeutic potential in cancer treatment. Recently, *B. proteolyticus* strain OSUB18, isolated from switchgrass, has been demonstrated to promote plant growth and regulate reactive oxygen species (ROS) production. Here, we investigated the anticancer potential of this plant-associated microbe and demonstrated that its crude extracts exhibited broad-spectrum cytostatic activity across diverse solid tumor cell lines. Importantly, OSUB18 extracts displayed limited cytotoxicity toward normal human mammary epithelial cells and mouse fibroblasts. This potentially selective cytostatic activity was retained within a water soluble, heat-labile, and protease-resistant fraction. The extracts primarily suppressed cell proliferation, accompanied by increased lipid peroxidation, with preliminary evidence suggesting the potential involvement of ferroptosis. These findings suggest that the bioactive component may be a selective cytostatic agent with multifaceted mechanisms of action.

## Background

Cancer remains the second leading cause of worldwide mortality and one of the most significant challenges to public health. According to 2020 estimates, nearly 10 million deaths (1 in 6) were attributed to cancer [1]. The landscape of cancer treatment has undergone substantial advances, beginning with conventional cancer therapeutic strategies such as chemotherapy, hormone therapy, radiation, and surgery. More recently, novel therapies have emerged, including targeted therapy, immune checkpoint inhibitors (blocking checkpoint proteins like PD-1, PD-L1, CTLA-4) and chimeric antigen receptor (CAR) T-cell therapy [2]. These advancements have dramatically improved outcomes in cancer patients [2]. Nonetheless, advanced malignancies, such as metastatic triple-negative breast cancer (TNBC), castration resistant prostate cancer (CRPC), and non-small cell lung cancer (NSCLC), continue to pose major challenges due to relapse, acquired resistance, a lack of response in a substantial subset of patients, and immune evasion. Many approaches are associated with irreversible off-target treatment-related toxicities, which further complicates long-term disease management [3]. Therefore, effective therapeutics that improve survival outcomes and the quality of life for patients with advanced cancer remain a critical need.

Nature boasts immense chemical diversity, with natural products serving as a rich source of cancer chemotherapeutics [4]. Historically, many of the most effective anticancer agents originate from natural sources. Doxorubicin, used extensively for treatment of a broad range of malignancies including breast cancer, lymphoma, and leukemia, is a derivative of daunorubicin from *Streptomyces* [5]. Mitomycin C, which has demonstrated efficacy in bladder, stomach, and pancreas cancer patients, is a natural product from *S. caespitosus* [6]. These examples underscore the enduring value of natural products for anticancer drug discovery and development.

Microorganisms remain an abundant and relatively underexplored reservoir of chemical and biological diversity with potentially useful activities. Among bacteria, *Bacillus* produces a wide range of bioactive secondary metabolites, including lipopeptides, polyketides, and bacteriocins [7]. *Bacillus* strains have also demonstrated selective cytotoxicity against cancer cells. Reported inhibitory components have included extracellular polymeric substances (EPS) from *B. subtilis* [8], surface layer protein and parasporins from *B. thuringiensis* [9, 10], lipopeptide surfactin derived from *B. subtilis* [11], and a heat-stable lysozyme derived from the thermophilic bacterium *B. paralicheniformis* BplzC [12]. Here we investigate the anticancer potential of *B. proteolyticus* strain OSUB18 against multiple cancer cell lines. Isolated from switchgrass, OSUB18 has been demonstrated to promote plant growth and regulate reactive oxygen species (ROS) production [13, 14]. As elevated concentrations of ROS can induce oxidative damage and trigger cell death, manipulation of ROS level presents a promising strategy for cancer therapy [15]. We observed that bioactive component of OSUB18 exhibits cytostatic effects on various human and mouse cancer cell lines while sparing normal epithelial cells and fibroblasts. Our findings demonstrate that OSUB18 produces a soluble, heat-sensitive, and protease-resistant component that selectively suppresses cancer cell proliferation while inducing lipid peroxidation.

## Methods

### Cell culture

The human and mouse breast cancer cell lines MDA-MB 231, 4T1 and E0771, human and mouse CRPC lines PC-3 and TRAMP-C1, human glioblastoma U251, human and mouse melanoma LM-MEL 53 and B16F10, human colon cancer cells HCT116, human NSCLC cell line A549, human normal mammary epithelial cell MCF10A and mouse fibroblasts 3T3 were obtained from American Type Culture Collection. MDA-MB 231, and 4T1 cells were maintained in Dulbecco′s Modified Eagle′s Medium/Nutrient Mixture F-12 Ham (DMEM/F12) (Sigma Aldrich, D8437) with 10% fetal bovine serum (FBS) (Sigma Aldrich, F2442), penicillin (100 units/mL) and streptomycin (0.1 mg/mL) (P/S) (Sigma Aldrich, P4333). TRAMP-C1 and 3T3 cells were maintained in Dulbecco’s Modified Eagle’s Medium (DMEM) (Sigma Aldrich, D6429) with 10% FBS and P/S. MCF10A cells were maintained in DMEM/F12 with 5% horse serum (Sigma Aldrich, 12449C) and P/S, 20 ng/mL EGF (PEPROTECH, AF-100-15), 0.5 mg/mL hydrocortisone (Sigma, H0888), 100 ng/mL Cholera Toxin (Sigma Aldrich, C8052), 10 μg/mL Insulin (Sigma Aldrich, I6634). E0771 cells were maintained in RPMI-1640 Medium (Sigma Aldrich, P3813) with 10% FBS, 10 mM HEPES (pH7.4, Sigma Aldrich, H4034) and P/S. PC-3, TRAMP-C1, U251, LM-MEL 53, B16F10, HCT116, and A549 cells were maintained in RPMI-1640 Medium (Sigma Aldrich, P3813) with 10% FBS and P/S.

To prepare 3D laminin-rich extracellular matrix on-top cultures, we trypsinized cells from tissue culture plastic and seeded single cells onto a thin gel of Growth Factor Reduced Basement Membrane Matrix (Matrigel, Corning, 354230), and addition of medium containing 5% Matrigel. Cancer cells were seeded at a density of 4,000 cells per well in a 24-well culture plate and maintained in their standard propagation media. Following treatment with microbial-derived extracts, the cell colonies in 3D were imaged, and their diameters were quantified using the NIS Elements AR software.

### Growth of bacterial strains and production of extracts

For initial screening experiments, bacterial extracellular exudates (BEE) were produced as previously described [16]. In general, the bacterial cultures were grown on Tryptic Soy Agar (TSA) plates at 28 °C for a period of two days. Subsequently, bacterial cells from each strain (with four plates each) were harvested by washing them off the TSA plates using 10 mL sterile water and collected into a 50 mL tube. For control purposes, 10 mL of sterile water was similarly used to wash an uninoculated TSA plate. The optical density at 600 nm (OD600) was measured in the vortexed solution to quantify the cell numbers of each bacterial culture. The bacterial suspension was centrifuged at 4 °C, 4,300 rpm for 30 min to obtain clarified supernatant. To ensure the supernatant was free of cells, 100 µL of each supernatant was plated onto a new TSA plate to confirm the absence of viable bacterial cells. Generally, each 10 mL of bacterial solution yielded 8 mL of 1× native BEE. The 1× native BEE was further concentrated to 10× native BEE using a freeze dryer.

### Fractionation of concentrated extracts

Extracts used in fractionation experiments were produced by inoculating OSUB18 into 20 ×150 mm test tubes containing 5 mL Tryptic Soy Broth (TSB) (17 g tryptone, 5 g soytone, 2.5 g glucose, 5 g NaCl, 2.5 g K_2_HPO_4_, 1 L distilled H_2_O). Cultures were grown on a roller drum set at 30 °C, 80 rpm. After 24 hours, a 1 mL aliquot of the seed culture was used to inoculate 500 mL Fernbach flasks containing 100 mL of TSB. Flask cultures were grown on a shaking incubator at 30 °C, 200 rpm. After 16 hours, cultures were harvested by centrifugation, and the clarified supernatant freeze dried. 700 mg of freeze-dried material from flask cultures was re-dissolved in 5 mL of sterile ultrapure water and centrifuged at 10,000 rpm. The residual pellet material was resuspended with 3 mL sterile ultrapure water and re-centrifuged two times. Clarified supernatants from the initial resuspension and pellet washes were applied onto a 50 kDa MWCO ultrafiltration concentrator and continually centrifuged at 10 °C, 3,300 rpm until no additional filtrate passed through. The resulting filtrate was then applied onto a 30 kDa MWCO ultrafiltration concentrator as above. The retentate from both concentrators, and the 30 kDa filtrate were frozen at −80 °C and lyophilized to dryness. Total protein levels in concentrated extracts and retentate were measured using Pierce™ BCA Protein Assay Kit (Thermo Fisher Scientific, 23227) using bovine serum albumin as a standard.

### Heat and protease treatment

The 50 kDa retentate was boiled for 10 minutes at 100 °C to determine the thermal stability of the bioactive component. For protease digestion, the retentate was adjusted to protein concentration of 1 mg/mL, then treated with 0.3 mg/mL trypsin or 0.1 mg/mL proteinase K at 37°C for 90 min. Phenyl-methyl sulfonyl fluoride (PMSF) in isopropanol was added (1 mM) to stop protease digestions [17].

### Cell viability assay

Cell viability assay was performed using the Cell Counting Kit-8 (CCK8, Sigma Aldrich, 96992). Cancer cells were seeded to a 96-well cell culture plate (10,000 cells per well). Control (the extract of uninoculated TSA plate) or microbial-derived bioactive extracts were added to the cell culture medium. After 3-day treatment, 10 μL of the CCK-8 solution was added to each well of the plate, according to the manual instructions. The plate was incubated for 1–3 h at 37 °C in the cell incubator. The 96-well plate was measured on a Tecan Sunrise Plate Reader (Tecan) at 450 nm. For each cell line, the absorbance values in the drug treatment group were normalized to the value in the control group.

### Cell proliferation assay

Cell proliferation was examined using Click-iT™ EdU Cell Proliferation Kit for Imaging, Alexa Fluor™ 488 dye (Invitrogen, C10337). Cancer cells were seeded into 8-well Cell Culture Chamber Slide (Alkali Scientific, TCC008). Control or microbial-derived bioactive extracts were added to the cell culture medium. After 2 days, 5-ethynyl-2-deoxyuridine (EdU) solution was added to the cell cultured medium at a final concentration of 10 μM and incubated for 8 h at 37 °C. After incubation, medium was removed from the cells and fixed with 4% PFA at room temperature for 15 min, followed by 0.5% Triton® X-100 permeabilization. Cells were incubated with the Click-iT® reaction cocktail at room temperature for 30 min to detect EdU-positive cells. Cells were incubated with DAPI solution for 2 min and then imaged using a Nikon Eclipse Ts2 microscope. EdU positive ratio was quantified by counting EdU and DAPI positive cells.

### Cell apoptosis assay

Cell apoptosis assay was performed using the Dead Cell Apoptosis Kits with Annexin V for Flow Cytometry (Invitrogen, V13242). In general, after treatment with control (the extract of uninoculated TSA plate) or microbial-derived bioactive extracts, cancer cells were harvested and incubated with FITC Annexin V and 1 μg/mL propidium iodide (PI). Following 15 min incubation at room temperature, stained cells were analyzed using an Attune NxT flow cytometer (Invitrogen). Data were analyzed by FlowJo software.

### Lipid peroxidation assay

Lipid peroxidation was measured by BODIPY™ 581/591 C11 staining (Invitrogen, D3861) of control or treated cancer cells. Cancer cells were seeded in 24-well culture plates. The next day, the cultured cancer cells were treated with control or microbial-derived bioactive extracts, with and without ferroptosis activator RSL3 (1 μM, MedChemExpress, HY-100218A) or ferroptosis inhibitor ferrostatin-1 (Fer-1, 5 μM, MedChemExpress, HY-100579) and grown for additional 2-3 days. To determine lipid peroxidation, cancer cells were incubated with 5 μM BODIPY™ 581/591 C11 (Invitrogen, D3861) for 30 min at 37 °C. Oxidized and non-oxidized C11 stained cells were imaged using Nikon Ts2R Eclipse Inverted Microscope.

### Transcript analysis of marker genes

Total RNA was extracted from control and treated cells using TRIzol reagent (Invitrogen, 15596018). cDNA was synthesized using the SuperScript™ IV First-Strand Synthesis System (Invitrogen, 18091050) from 2.0 μg of RNA for each sample according to the manufacturer’s instructions. Reverse transcription quantitative polymerase chain reactions (RT-qPCR) were carried out with SYBR green PCR master mix reagents using a QuantStudio™ 3 Real-Time PCR System (Thermo Fisher Scientific, A28567). Thermal cycling was performed as previously described [18]: an initial 95°C for 30 sec, followed by 40 cycles of amplification at 95°C for 5 sec, 55°C for 30 sec, and 72°C for 15 sec. The relative quantification of gene expression for each sample was analyzed by the △ threshold cycle (△△CT) method. Primers used for amplification cell death related genes and 18S rRNA are listed in Table 1.

**Table 1.**
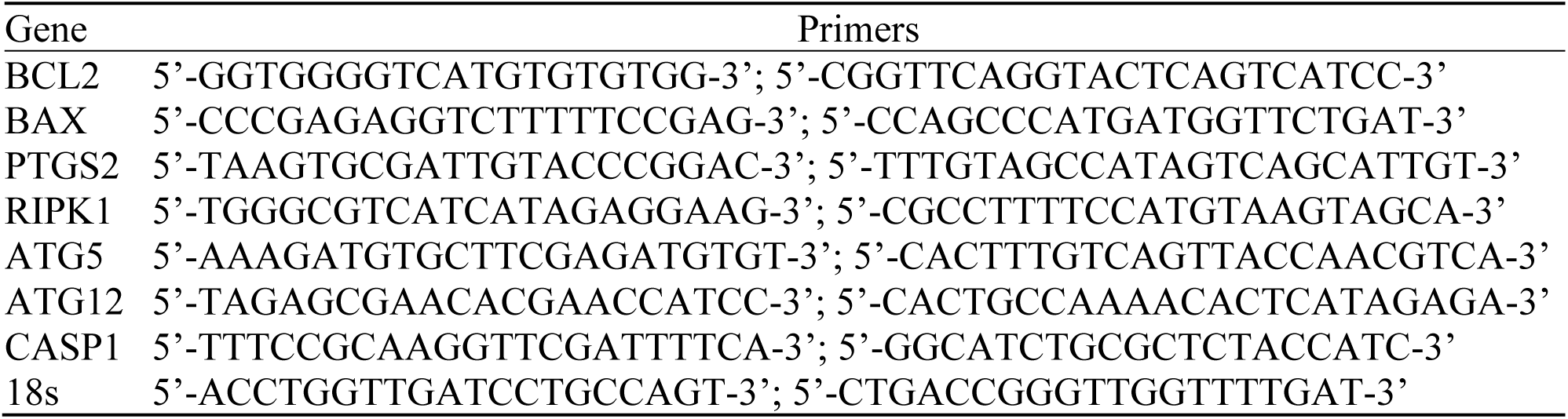
Primers used in RT-qPCR.

### Statistical Analysis

Primary experiments were repeated at least three biological replicates; certain preliminary mechanistic screening were conducted as exploratory assessments. Data are presented as mean ± standard error of the mean (SEM). Statistical analyses were performed using GraphPad Prism software. For comparisons between two groups, independent Student’s T-test was used. For comparisons among multiple groups, one-way analysis of variance (ANOVA) followed by post hoc tests was applied, as specified in the figure legends.

## Results

### Broad-spectrum cytostatic activity of OSUB18 crude extracts

We treated multiple cancer cell lines (including MDA-MB 231, 4T1, E0771, PC-3, TRAMP-C1, U251, LM-MEL 53, B16F10, A549, and HCT116 cells) with crude extracts of switchgrass-derived *Bacillus* isolates [14] and monitored cell growth in two-dimensional (2D) plastic culture. After three days of treatment, cell viability was measured using the CCK8 assay [18]. Extracts from strain OSUB18 significantly inhibited the growth of various solid cancer cell lines, with limited effect on the growth of normal mammary epithelial MCF10A cells and 3T3 mouse fibroblasts (Figure 1A, 1B). Exploratory observations suggested a concentration-dependent reduction in cell growth (Supplemental Figure 1). These findings indicate that the extracts have broad-spectrum cytostatic effects on multiple cancer cell lines, with limited cytotoxicity toward normal cells.

**Figure 1.**
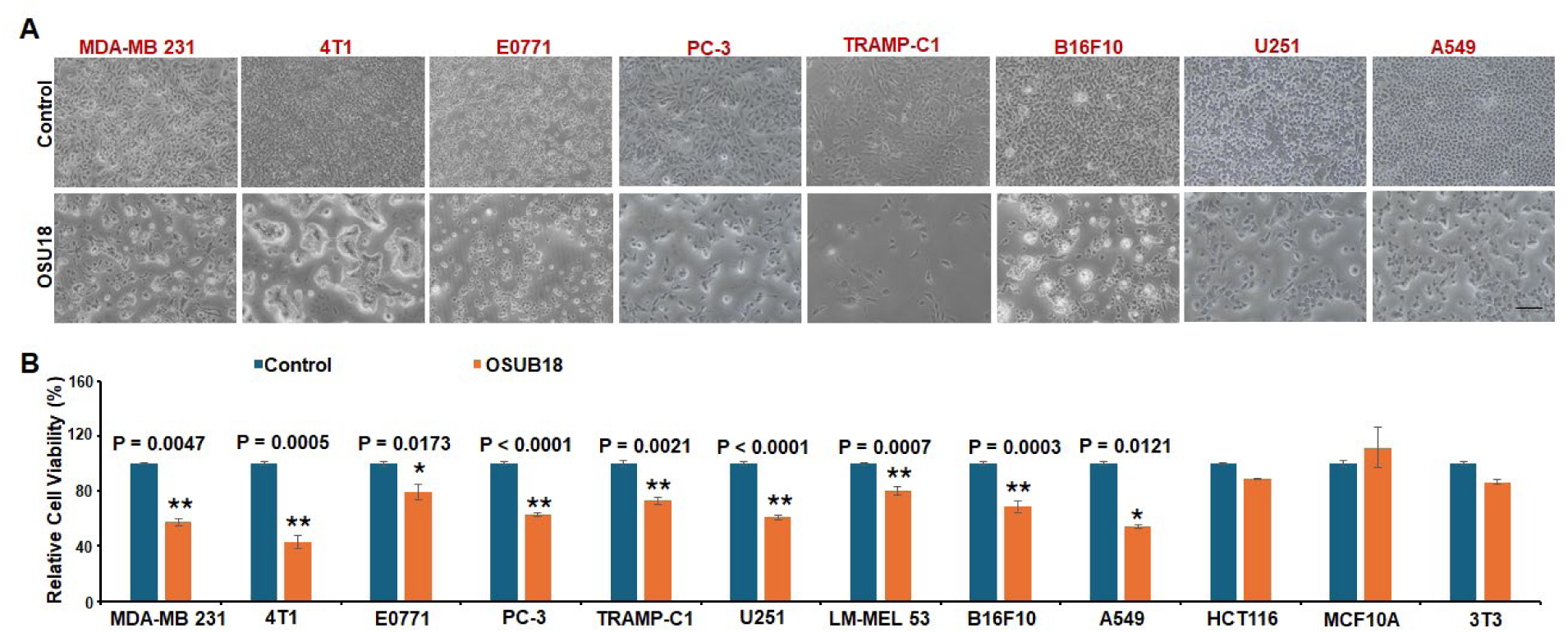
Extracts derived from *B. proteolyticus* strain OSUB18 exhibit cytostatic efficacy. (A) (B) Cell images and cell viability of various cancer cell lines and normal mammary epithelial MCF10A and normal fibroblasts 3T3 cells treated with 10 μL control or crude extracts (∼50 mg/mL) from *Bacillus proteolyticus* strain OSUB18 for 3 days in 96-well plate (100 μL culture medium each well), measured by CCK8 assay. Scale bar: 200 μm. Results are presented as mean ± SEM, n = 3, independent Student’s t test. Control: the extract of uninoculated TSA plate. OSUB18: extracts from OSUB18.

### Suppression of cancer growth in 3D models

Due to low cost, ease of use, and suitability for high-throughput screening, 2D monolayer cell culture remains the primary method for evaluating anticancer drug efficacy. However, cancer cells grown on stiff plastic surfaces exhibit limited cell–cell interactions and undergo significant physiological and molecular alterations that fail to accurately recapitulate the tumor microenvironment [19, 20]. In contrast, three-dimensional (3D) cell cultures more closely mimic the complex tumor niche, providing a superior approach for investigating tumor biology and drug response [19, 20]. Therefore, we employed a 3D Matrigel culture model to further assess the cytostatic efficacy of OSUB18 crude extracts. Cancer cells were seeded into Matrigel and treated with either OSUB18 extracts or control. After three days of incubation, cancer cell colonies were imaged and their sizes were measured. Consistent with our 2D culture findings, our results showed that cancer cell colonies (4T1, U251, PC-3, and A549) grown in 3D Matrigel were markedly smaller in the extract-treated groups compared to the control groups (Figure 2A, 2B). Thus, OSUB18-derived extracts also suppressed cancer growth within a 3D Matrigel matrix.

**Figure 2.**
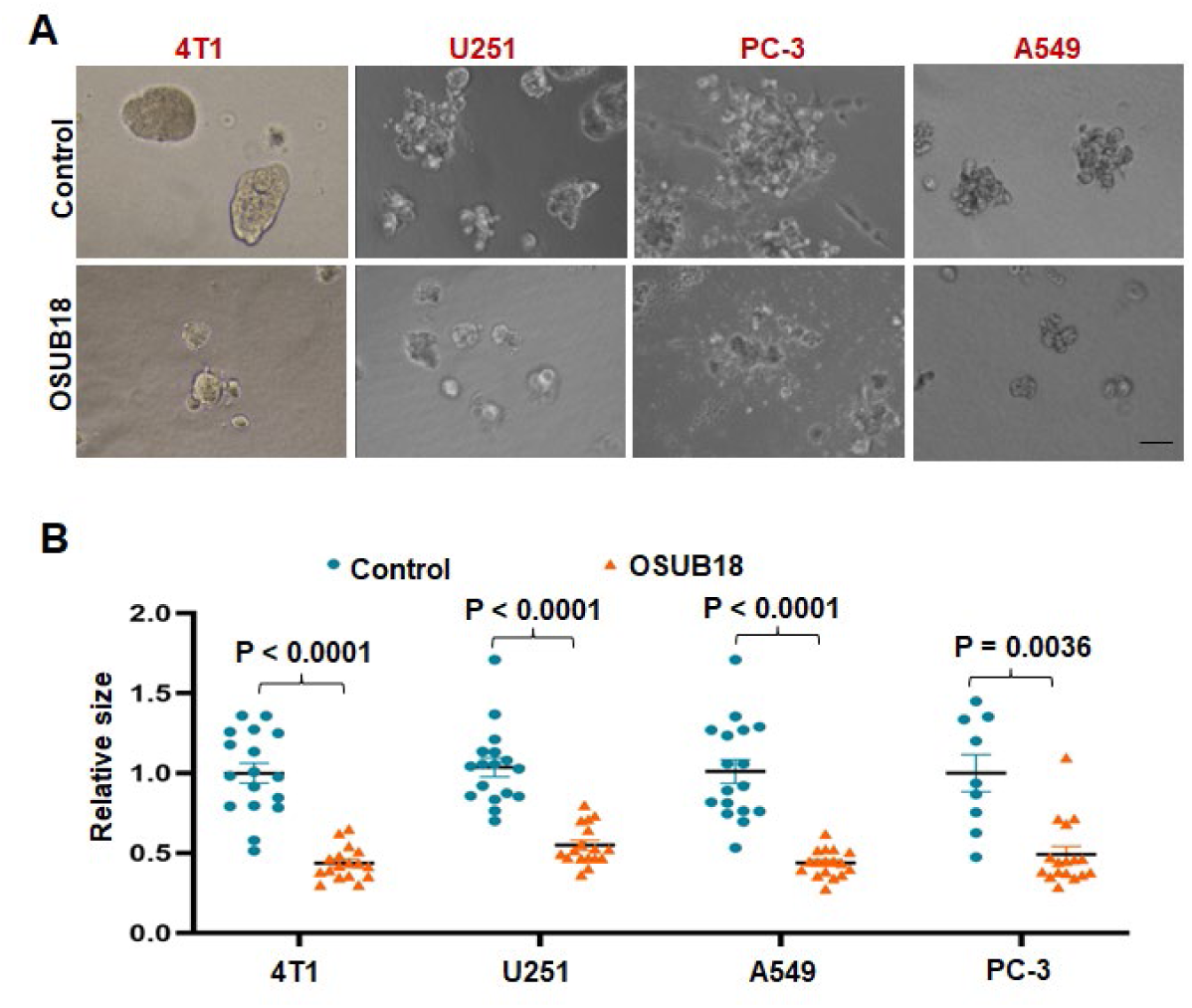
Assessment of cytostatic efficacy of OSUB18 extracts in 3D model. (A) Images of 4T1, U251, PC-3 and A549 cells cultured in 3D Matrigel, treated with 50 μL control or OSUB18-derived extracts in 500 μL culture medium for 3 days in a 24-well culture plate. Scale bar: 50 μm. (B) Clone diameters of 4T1, U251, PC-3, and A549 cells cultured in 3D Matrigel, treated with control or OSUB18-derived extracts. Results are presented as mean ± SEM, n = 3 (randomly selected images for each group), independent Student’s t test. Control: the extract of uninoculated TSA plate. OSUB18: extracts from OSUB18.

### Cancer cell proliferation

We performed various cellular assays to examine the underlying mechanisms of action of the crude extracts. Since many anticancer agents exert their effects by inhibiting cancer cell proliferation and/or inducing cell death, we first assessed whether the extracts alter cancer cell proliferation. The EdU incorporation assay [21] was performed to examine if the extracts may perturb DNA synthesis. Treatment with the extracts significantly reduced the proportion of EdU-positive cells across multiple cell lines (MDA-MB 231, 4T1, A549, B16F10, LM-MEL 53, TRAMP-C1, and U251) compared with control groups (Figure 3A-3C). These results confirm that the crude extracts exert a potent anti-proliferative effect across diver tumor types.

**Figure 3.**
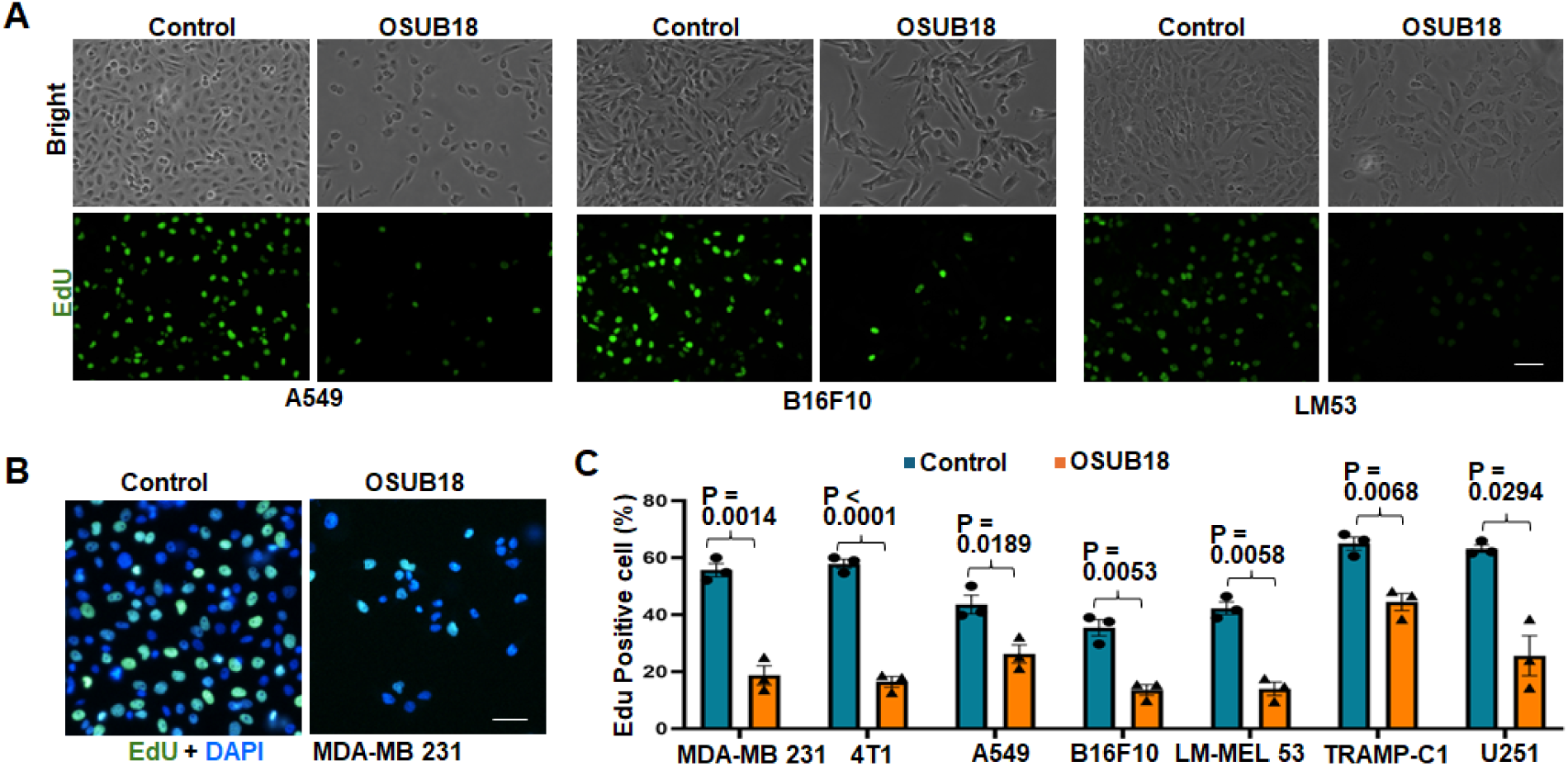
OSUB18 extracts inhibit cell proliferation in cancer cells. (A) Representative bright and EdU staining images of cancer cells treated with 20 μL control or OSUB18-derived extracts for 3 days in 8-chamber cell culture slide (200 μL culture medium). (B) Representative confocal images of DAPI and EdU staining in TNBC cells treated with 20 μL control or OSUB18-derived extracts in an 8-chamber for 3 days. (C) Quantification of cell proliferation in MDA-MB 231, 4T1, A549, B16F10, LM-MEL 53, TRAMP-C1 and U251 cells treated with control or OSUB18-derived extracts. Results are presented as mean ± SEM, n = 3 (randomly selected images for each group), independent Student’s t test. Scale bar: 50 μm.

### Lipid peroxidation

Many conventional anticancer drugs, such as doxorubicin, cisplatin, and paclitaxel, induce cancer cell death primarily through apoptosis, a programmed cell death mechanism critical for eliminating damaged or abnormal cells [22–24]. To determine whether the extracts also induce apoptosis, we analyzed control and extract-treated cells using Annexin V/PI flow cytometry. Our preliminary analyses indicated that treatment with the extracts did not increase in the population of apoptotic cells. Instead, a trend toward a reduced percentage of early apoptotic cancer cells was observed in the extract-treated groups compared to controls, whereas late apoptotic cell populations appeared comparable between groups (Supplemental Figure 2A, 2B). However, microscopic examination revealed an increased number of detached or severely damaged cells following extract treatment. It is possible that this nonviable population was underrepresented in the flow cytometry analysis due to cell loss during sample preparation or exclusion during gating. Therefore, additional measurements of cell death are needed to comprehensively determine the effects of extract treatment on cell death pathways.

We performed RT-qPCR analysis of genes associated with multiple cell death pathways, including ferroptosis (Prostaglandin-Endoperoxide Synthase 2, *PTGS2*) [25], apoptosis (B-cell CLL/lymphoma 2, *BCL2* and BCL2 Associated X, *BAX*), necroptosis (Receptor Interacting Serine/Threonine Kinase 1, *RIPK1*), autophagy (Autophagy protein 5 and 12, *ATG5, ATG12*), and pyroptosis (Caspase 1, *CASP1*) [26], to explore their involvement with the observed inhibition. Our exploratory data showed that transcripts were increased for *PTGS2* in extract-treated cancer cells compared to control groups. Conversely, no substantial differences in transcript abundance was observed for genes associated with apoptosis, autophagy, or pyroptosis (Figure 4A, Supplemental Figure 2C).

**Figure 4.**
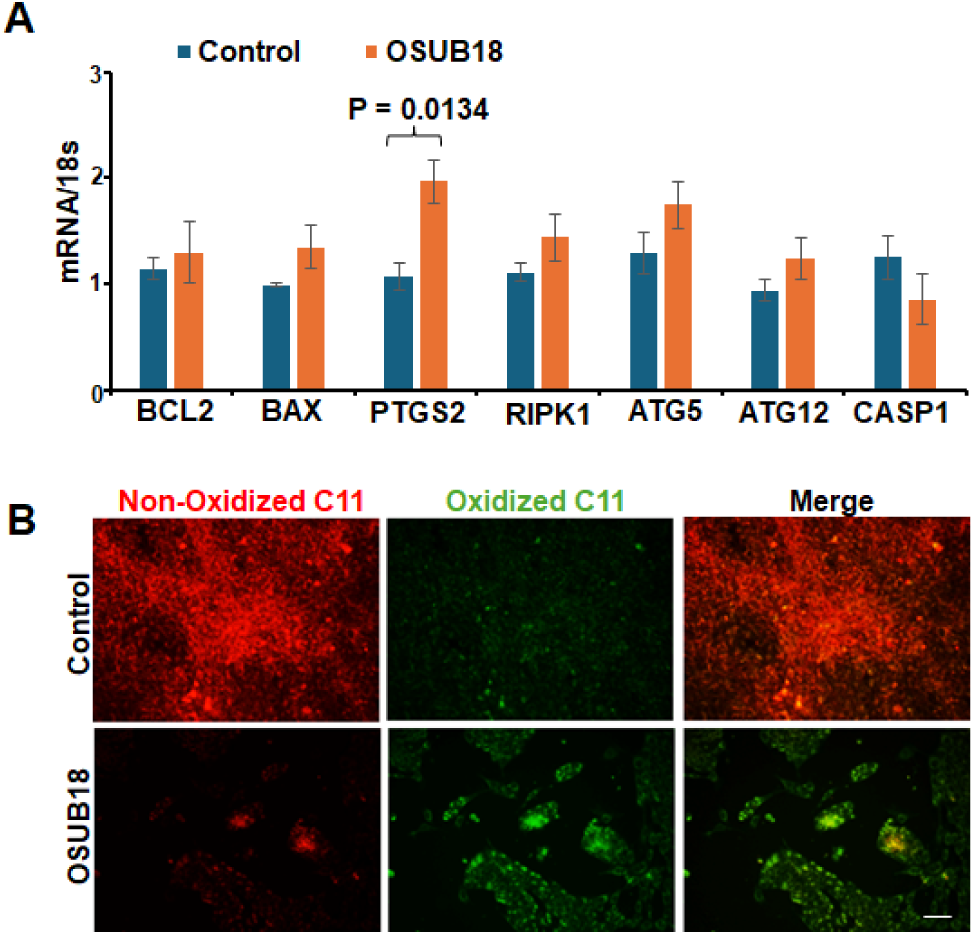
Increased lipid peroxidation in extract-treated cancer cells. (A) RT-qPCR analysis of cell death related genes in MDA-MB 231 cells treated with 200 μL control or OSUB18-derived extracts (2 mL culture medium in 60 mm culture dish) for 3 days. Results are presented as mean ± SEM, n = 2, independent Student’s t test. (B) BODIPY™ 581/591 C11 (5 μM) staining in 4T1 cells treated with 50 μL control or OSUB18 extracts (500 μL culture medium in 24-well culture plate) for 3 days. Scale bar: 50 μm. Control: the extract of uninoculated TSA plate. OSUB18: extracts from OSUB18.

Lipid peroxidation arises from the oxidation of membrane lipids by ROS and serves as both a marker and a driving mechanism for oxidative stress-induced cell death pathways, including ferroptosis [27]. To examine the potential involvement of ferroptosis, we measured lipid peroxidation using BODIPY™ 581/591 C11 staining, a probe commonly used to detect lipid peroxidation in live cultured cells [28]. Fluorescence imaging revealed that extract-treated TNBC cells displayed an enhanced signal for oxidized C11, whereas control cells exhibited the minimal baseline signal (Figure 4B). These preliminary findings indicate enhanced lipid peroxidation following extract-treatment, suggesting potential involvement of ferroptosis-related pathways.

Additional validation of ferroptosis markers and identification of their downstream targets are required to fully establish the mechanisms of action.

### Activity of fractionated extracts

Crude extracts were fractionated using ultrafiltration devices (Figure 5A). Cell viability assays revealed that the inhibitory activity resided predominantly within the >50 kDa retentate fraction (Figure 5A). To determine whether the bioactive component in this fraction is sensitive to heat treatment, the retentate was boiled for 10 minutes. This treatment resulted in a significant loss of inhibitory activity (Figure 5B), indicating that the bioactive component is heat sensitive. Interestingly, digestion with trypsin or proteinase K [17], did not diminish its inhibitory activity (Figure 5C). These results suggest the extracts contain a water soluble, heat-sensitive, and protease-resistant component enriched within the >50 kDa retentate, although its exact identity remains to be determined.

**Figure 5.**
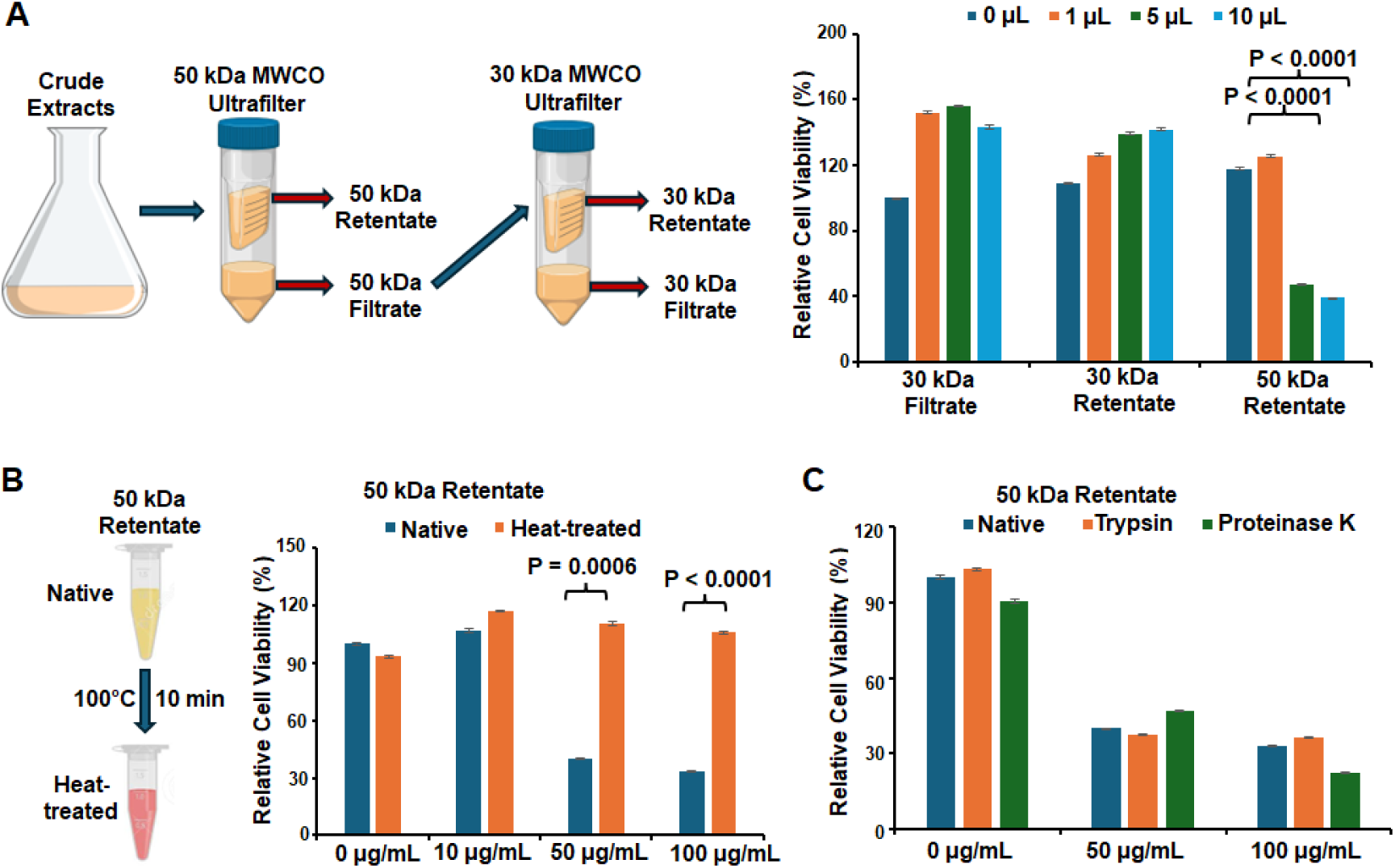
The bioactive component of OSUB18 extracts is heat-sensitive and protease-resistant. (A) Cell viability of 4T1 treated with different volume (0 μL, 1 μL, 5 μL, 10 μL) of filtrate and retentate of OSUB18 crude extracts separated by 30 kDa and 50 kDa MWCO Ultrafilter in 96-well plate (100 μL culture medium) for 3 days, measured by CCK8 assay. Results are presented as mean ± SEM, n = 3, one-way ANOVA followed by Dunnett’s multiple comparisons test. (B) Cell viability of 4T1 treated with diffenent concentrations (0 μg/mL, 10 μg/mL, 50 μg/mL, 100 μg/mL) of native or heat-treated (100°C for 10 min) OSUB18 50 kDa retentate in 96-well plate for 3 days, measured by CCK8 assay. Results are presented as mean ± SEM, n = 3, independent Student’s t test. (C) Cell viability of 4T1 treated with different concentrations (0 μg/mL, 50 μg/mL, 100 μg/mL) of native or digested OSUB18 50 kDa retentate in 96-well plate for 3 days. Retentate was pre-treated with trypsin or proteinase K at 37°C for 90 min. Results are presented as mean ± SEM, n = 3.

Next, we examined the effects of the *>*50 kDa retentate on cell proliferation and lipid peroxidation. Consistent with our findings using crude extracts, EdU staining results demonstrated that the retentate significantly inhibited cancer cell proliferation and overall growth (Figure 6A, 6B). To determine whether the *>*50 kDa retentate fraction induces lipid peroxidation in cancer cells, TNBC cells were treated with retentate in the presence or absence of ferrostatin-1 (Fer-1), a widely used ferroptosis inhibitor that prevents lipid peroxidation [29]. The addition of Fer-1 partially rescued the cytostatic activity of the *>*50 kDa retentate (Figure 6B). Heat treatment completely abolished the cytostatic effect of the retentate (Figure 6B). Fluorescence imaging confirmed that the *>*50 kDa retentate increased the oxidized C11 staining signal in TNBC cells (Figure 6C, 6D). This increase was successfully abrogated by either Fer-1 co-treatment or by using the heat-treated retentate (Figure 6C, 6D). These findings suggest that the cytostatic effect of this fraction is partially ferroptosis-dependent, with additional mechanisms likely contributing to the observed reduction in cell viability.

**Figure 6.**
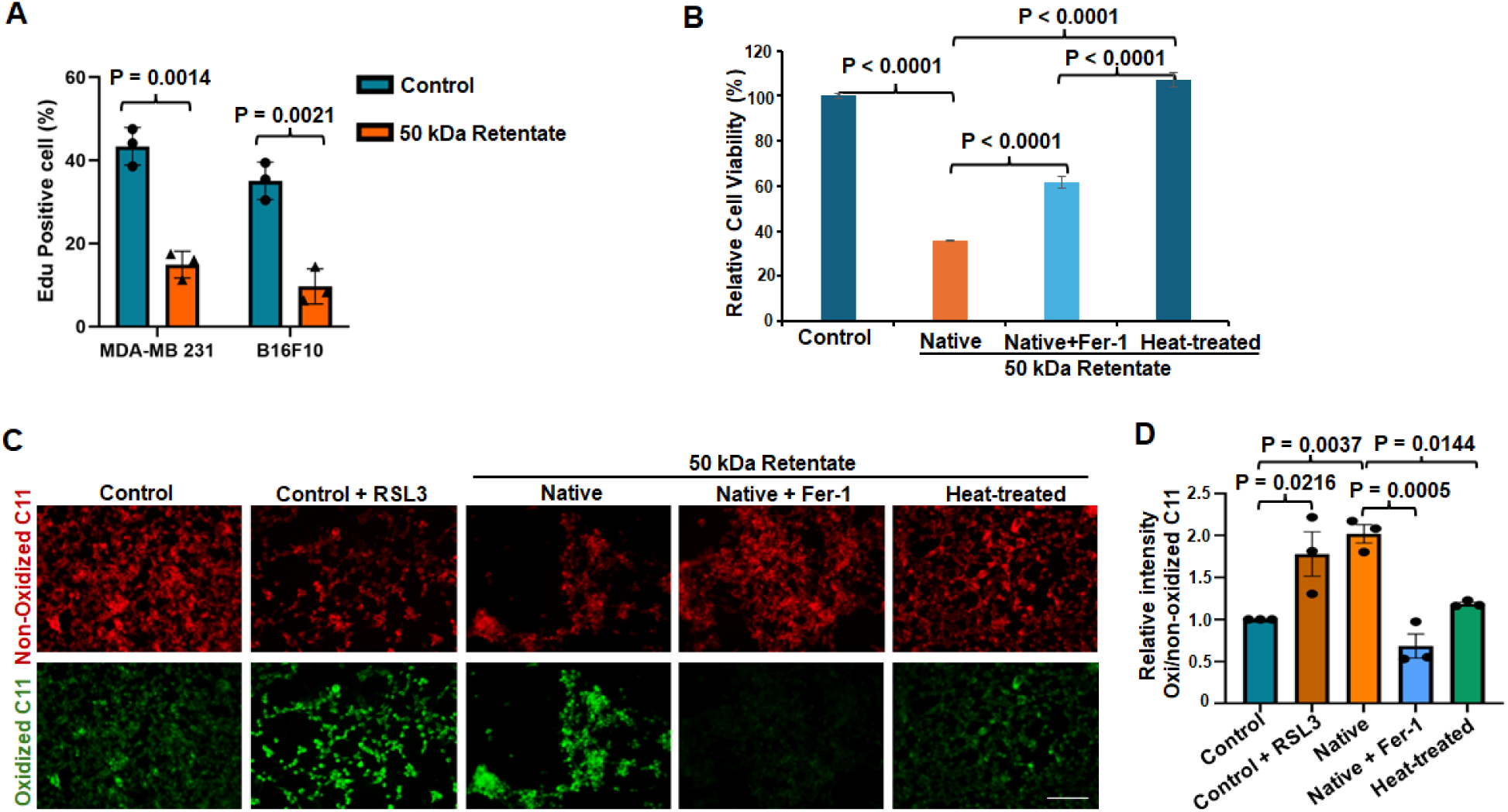
The 50 kDa retentate fraction suppresses cancer cell growth. (A) Quantification of cell proliferation (EdU incorporation) in MDA-MB 231 and B16F10 treated control (30 kDa retentate) or 50 kDa retentate in 8-chamber cell culture slide for 3 days. Results are presented as mean ± SEM, n = 3, independent Student’s t test. (B) Cell viability of 4T1 treated 30 μg/mL control (30 kDa retentate) or 50 kDa retentate combined with DMSO or Fer-1 in 96-well plate for 3 days. Results are presented as mean ± SEM, n = 3, one-way ANOVA followed by Turkey’s multiple comparisons test. (C), (D) BODIPY™ 581/591 C11 staining images (C) and quantification (D) in 4T1 cells treated with 30 μg/mL control (30 kDa retentate) or 50 kDa retentate combined with DMSO, RAL or Fer-1 in 24-well culture plate for 3 days. RSL3: cells treated with 1 μM ferroptosis inducer RSL3 as positive control. Fer-1: cells treated with 5 μM ferrostatin-1. Native, Heat-treated: original or boiled 50 kDa retentate of OSUB18 extracts. Scale bar: 50 μm. Results are presented as mean ± SEM, n = 3, one-way ANOVA followed by Turkey’s multiple comparisons test.

Consistent with the crude extracts, the fractionated retentate exhibited a cytostatic effect by inhibiting cancer cell proliferation and increasing lipid peroxidation. Alleviation of this cytostatic effect by Fer-1 suggests that the activity may partially be attributed to ferroptosis.

## Discussion

Despite significant advancements in cancer therapy over the past few decades, advanced malignancies persist as a major challenge. Cancer remains the second leading cause of mortality worldwide. Thus, new treatment approaches that minimize toxicity and prolong patient survival are urgently needed. In this study, we demonstrated crude extracts of *B. proteolyticus* strain OSUB18 exhibits broad-spectrum and potentially selective cytostatic activity across various solid tumor cell lines, including those of breast, lung, prostate, colorectal, brain and skin, while sparing normal epithelial cells and fibroblasts. The extracts appear to suppress cell proliferation, with an accompanied increased lipid peroxidation. These cytostatic effects may be partially ferroptosis-dependent, with additional mechanisms likely contributing to the overall reduction in cell viability. Further characterization and exploration of the alternative pathways will be a focus of future studies.

The majority of conventional anticancer drugs, such as doxorubicin, cisplatin, and paclitaxel, are reported to induce cancer cell death primarily through DNA damage and apoptosis [22–24]. Conversely, only limited number of drugs and natural products, including cisplatin [30], curcumin [31], gallic acid [32], and acevaltrate [33], have been reported to exert anticancer activity by targeting glutathione peroxidase 4 (GPX4) and inducing ferroptosis. Previous studies have shown that crude extracts of OSUB18 influence cellular redox homeostasis in plants, altering ROS production and regulating ROS-associated gene respiratory burst oxidase homolog D (*RBOHD*, which encodes a NADPH oxidase) [14]. Given that ROS are key triggers or drivers of lipid peroxidation [34], the elevated lipid peroxidation observed in extract-treated cancer cells also suggests the possible involvement of ROS-mediated, ferroptosis-associated mechanisms.

Our results are similar to other reports of *Bacillus*-derived extracellular fractions exerting selective cytotoxic effects on cancer cells. These effects are frequently linked to oxidative stress-mediated pathways, characterized by intracellular ROS accumulation, loss of redox homeostasis and lipid peroxidation. Examples include EPS, derived from *B. subtilis* that exhibits strong ROS scavenging capacity [8]. Similarly, surfactin from *B. subtilis* has been reported to induce apoptosis through a ROS-regulated mitochondrial pathway [11], and BplzC from the thermophilic bacterium *B. paralicheniformis* exhibits selective cytotoxicity on cancer cells and ROS scavenging activity [12].

Ferroptosis represents an intriguing pathway for anti-tumor strategies due to its multifaceted roles in malignant tumors. Beyond disrupting cellular anion homeostasis, it has been shown to enhance T cells-mediated anticancer immune response and augments the efficacy of immunotherapy [35]. In out study, additional experiments, such as the genetic perturbation of key ferroptosis regulators (e.g., *GPX4*) and orthogonal biochemical assays (e.g., ROS measurements and iron dependency assays) [36], will be required to establish whether ferroptosis is a driver of the observed activity for OSUB18. Delineation of the molecular pathways by which the bioactive component modulates ROS homeostasis and lipid peroxidation signaling is the focus of our future investigations.

In recent years, microbial cytotoxic proteins have gained attention as a class of natural products with multivalent anticancer properties. Notable examples include azurin from *P. aeruginosa*, motility associated killing factor A from *V. cholerae*, surface layer protein from *B. thuringiensis*, and parasporins, a family of non-insecticidal proteins from *B. thuringiensis* that selectively kill cancer cells [9, 10, 37, 38]. These microbial cytotoxic proteins typically kill cancer cells by disrupting cancer cell membranes (pore-forming on cell membrane), inhibiting proliferation, inducing apoptosis, or in some cases, stimulating innate immune response [9, 10, 37, 38]. Remarkably, P28, a 28-amino-acid cell-penetrating peptide derived from azurin, has received orphan drug designation from the FDA for clinical trials in adult and pediatric cancer patients, underscoring the translational relevance of microbial molecules [39, 40]. Our results indicate that the potentially selective cytostatic activity of OSUB18 extracts is mediated by a soluble, heat-sensitive, and protease-resistant component. This may be a small molecule, soluble protein, or combinations thereof. To elucidate the molecular basis of its cytostatic effects, our future studies will focus on the purification and structure characterization of the bioactive component.

Our observations indicate that OSUB18 extracts have limited cytotoxicity toward two non-malignant cell lines. Following isolation of the bioactive component, a more comprehensive assessment across a broader panel of normal cell types, rigorous IC50 determination, cross-cell line dose standardization, and *in vivo* safety and toxicity studies will be required before considering its suitability as a therapeutic.

### Conclusion

We demonstrated that the crude extracts from *B. proteolyticus* OSUB18, a strain previously reported to promote plant growth and regulate ROS, exhibit broad-spectrum and potentially selective cytostatic activity across various solid tumor cell lines. The water soluble, heat-sensitive, and protease-resistant bioactive fraction suppresses cancer cell proliferation, accompanied by increased lipid peroxidation. These initial observations suggest the possible contribution of ferroptosis-related mechanisms, providing early insights into the anticancer potential of this plant-associated microbe.

## Availability of data and materials

All data generated or analyzed during this study are included in this published article.

## Competing interests

The authors declare no conflicts of interest.

## Funding

This research was funded by the start-up funding from The OSU Comprehensive Cancer Center to G.X.

## Author Contributions

Conceptualization, Y.X., G.X. and K.-S.J.; methodology, Y.X., G.X. and K.-S.J.; validation, G.X., B.K. and D.H.; investigation, B.K., D.H., P.Y. D.M. and G.X.; data curation, B.K. D.H. and G.X.; writing—original draft preparation, G.X.; writing—review and editing, K.-S.J. and Y.X.; supervision, G.X. All authors have read and agreed to the published version of the manuscript.

## Acknowledgments

PC-3 cell line is a kind gift from Dr. Dan Shu (The Ohio State University); U251, LM-MEL 53, B16F10 and A549 cell lines are kind gifts from Dr. Haichang Li (The Ohio State University).

## List of abbreviations

The following abbreviations are used in this manuscript:

2D: Two-dimensional
3D: Three-dimensional
Koda: Kilodalton
CAR: Chimeric antigen receptor
TNBC: Triple-negative breast cancer
CRPC: Castration resistant prostate cancer
NSCLC: Non-small cell lung cancer
DMEM/F12: Dulbecco’s Modified Eagle’s Medium/Nutrient Mixture F-12 Ham
DMEM: Dulbecco’s Modified Eagle’s Medium
FBS: Fetal bovine serum
P/S: Penicillin and Streptomycin
TSA: Tryptic Soy Agar
TSB: Tryptic Soy Broth
BEE: Bacterial extracellular exudate
RPM: Round Per Minute
CCK8: Cell Counting Kit-8
RT-qPCR: Reverse transcription quantitative polymerase chain reaction
Fer-1: ferrostatin-1
EdU: 5-Ethynyl-2-deoxyuridine
PTGS2: Prostaglandin-Endoperoxide Synthase 2
BCL2: B-cell CLL/lymphoma 2
BAX: BCL2 Associated X
RIPK1: Receptor Interacting Serine/Threonine Kinase 1
ATG5/12: Autophagy protein 5/12
CASP1: Caspase 1
RBOHD: Respiratory burst oxidase homolog D
GPX4: Glutathione peroxidase 4

## Notes

### Competing Interest Statement

The authors have declared no competing interest.

