## Supplemental figures for "Extracts of *Bacillus proteolyticus* strain OSUB18 exhibit cytostatic activity"

**Supplemental Information**

Supplemental Figure 1:

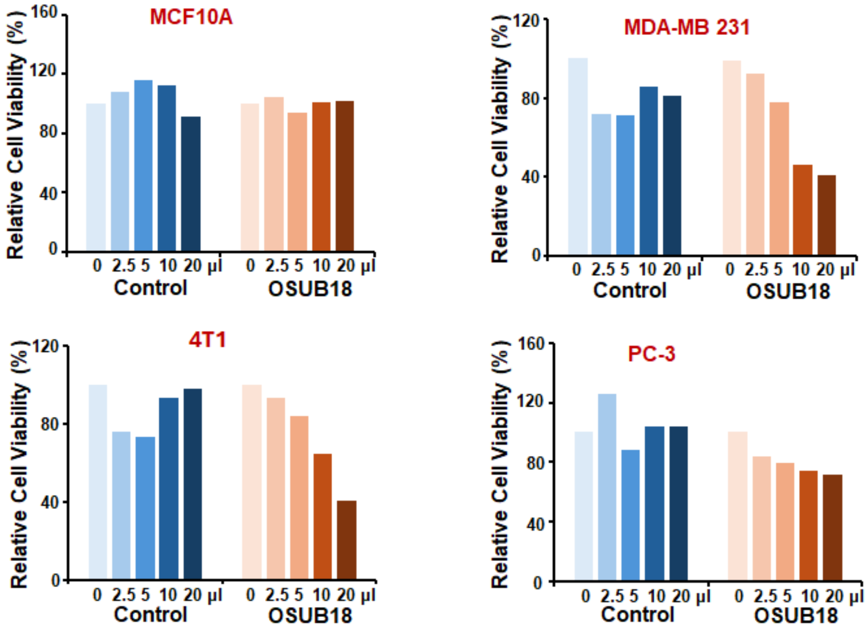


Supplemental Figure 1: Relative cell viability of cancer cell lines and normal epithelial cell treated with different doses of control or OSUB18-derived extracts in 96-well culture plate for 3 days.

Supplemental Figure 2:

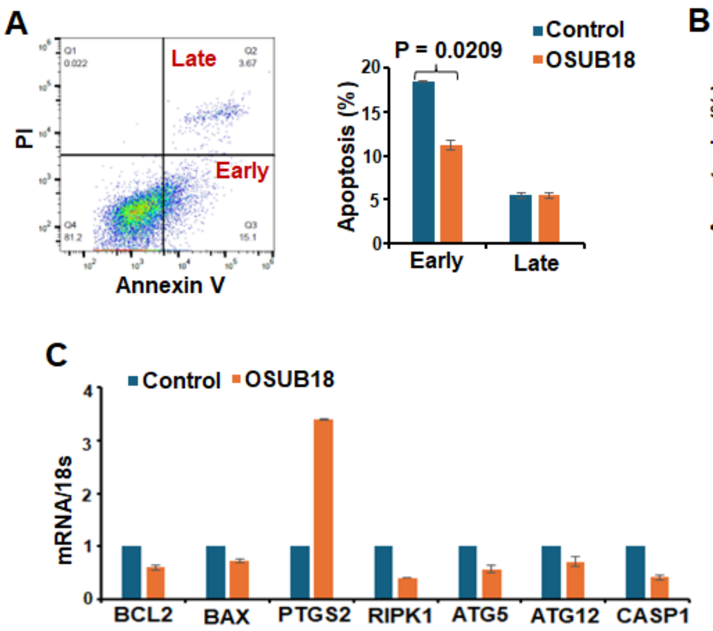


Supplemental Figure 2: (A) Annexin V and PI flow cytometry analysis cell apoptosis in U251 cancer cells treated with 200 µL control or OSUB18-derived extracts (2 mL culture medium in 60 mm culture dish) for 3 days. Results are presented as mean ± SEM, n = 3, independent Student's t test. (B) Annexin V & PI flow cytometry analysis cell apoptosis in 4T1 cells treated with 200 µL control or OSUB18-derived extracts (2 mL culture medium in 60 mm culture dish) for 3 days. Results are presented as mean ± SEM. (C) RT-qPCR analysis of cell death related genes in LM-MEL 53 cells treated with 200 µL control or OSUB18-derived extracts (2 mL culture medium in 60 mm culture dish) for 3 days. Results are presented as mean ± SEM, n = 2.
